# Sexual stage-induced long noncoding RNAs in the filamentous fungus *Fusarium graminearum*

**DOI:** 10.1101/270298

**Authors:** Wonyong Kim, Cristina Miguel-Rojas, Jie Wang, Jeffrey P Townsend, Frances Trail

## Abstract

Long noncoding RNA (lncRNA) plays important roles in morphological differentiation and development in eukaryotes. In filamentous fungi, however, little is known about lncRNAs and their roles in sexual development. Here we describe sexual stage-induced lncRNAs during the formation of perithecia, the sexual fruiting bodies of *Fusarium graminearum*. We have identified 547 lncRNAs whose expression was developmental stage-specific, with about 40% of which peaked during the development of asci, the sac-like structures containing meiospores. A large fraction of the lncRNAs were found to be antisense to mRNAs, forming 300 sense–antisense pairs. Although small RNAs (sRNAs) were produced from these overlapping loci, most of the antisense lncRNAs appeared not to be involved in gene silencing pathways. Genome-wide analysis of sRNA clusters identified many silenced loci at the meiotic stage. However, we found transcriptionally-active sRNA clusters, many of which were associated with lncRNAs. Also, we observed that many antisense lncRNAs and their respective sense transcripts were induced in parallel as the perithecia matured. To identify regulatory components for lncRNA expression, we analyzed mutants defective in the nonsense-mediated decay (NMD) pathway. A subset of the lncRNAs appeared to be targeted by the NMD before the perithecia formation, suggesting a suppressive role of the NMD in lncRNA expression during vegetative stage. This research provides fundamental genomic resources that will spur further investigations on developmental lncRNAs that may play important roles in shaping the fungal fruiting bodies.

## INTRODUCTION

Genomes of eukaryotes—from simple yeast to animals—are pervasively transcribed from noncoding intergenic regions and in antisense orientation from genic regions (Guttman et al. 2009; Xu et al. 2009). Long noncoding RNAs (lncRNAs) are loosely defined as noncoding transcripts longer than 200 nucleotides, which are mostly transcribed by RNA polymerase II and share common features with mRNAs other than protein-coding capacity (Kapranov et al. 2007). lncRNAs are versatile molecules that not only regulate gene expression, but also affect enzymatic activities and chromosome conformation (reviewed in Rinn and Chang 2012; Quinn and Chang 2015). Since the discovery of the *XIST* lncRNA required for X chromosome inactivation (Brockdorff et al. 1992), the roles of lncRNAs in developmental processes such as embryogenesis and tissue differentiation have been extensively studied in animals along with the advent of RNA-seq technologies (Fatica and Bozzoni 2014; Flynn and Chang 2014). Yet the full scope of the developmental roles of lncRNAs is far from understood.

In the highly divergent yeasts *Saccharomyces cerevisiae* (budding yeast) and *Schizosaccharomyces pombe* (fission yeast), the onset of sexual sporulation and the following meiotic divisions are tightly regulated by elaborate mechanisms involving lncRNAs (reviewed in Hiriart and Verdel 2013). In budding yeast, a promoter-derived lncRNA suppresses the expression of *IME1* (<u>i</u>nducer of <u>me</u>iosis 1), the master regulator for the sexual sporulation, by inducing heterochromatin formation in the promoter region of *IME1* during vegetative growth (van Werven et al. 2012). In addition, the transcription of another lncRNA antisense to *IME4* gene inhibits the expression of *IME4* by antagonizing the sense transcription (Hongay et al. 2006; Gelfand et al. 2011). Although there is no such conserved or analogous regulatory mechanism in fission yeast, lncRNAs also play diverse roles in the sexual sporulation, for example, sequestering RNA elimination factors that repress meiotic gene expression (Harigaya et al. 2006; Hiriart et al. 2012; Yamashita et al. 2012), and contributing homologous chromosome pairing (Ding et al. 2012). Despite the growing evidence of the regulatory roles in yeasts, information on lncRNA expression and function during fruiting body formation in filamentous fungi is scarce.

RNA quality control mechanisms are crucial for the regulation of lncRNA expression in budding yeast. The nuclear exosome is engaged in RNA processing and degradation of transcripts including lncRNAs that are specifically expressed during sexual sporulation; the deletion of *RRP6* encoding the exosome-associated exonuclease resulted in the accumulation of noncoding transcripts that otherwise remained silenced during vegetative growth (Davis and Ares 2006; Camblong et al. 2007; Lardenois et al. 2011). In human cells, promoter-derived transcripts were also ectopically expressed upon deletion of the exosome components including the homologous *RRP6*, suggesting the conserved role of the exosome for lncRNA expression in diverse eukaryotes (Preker et al. 2008). The nonsense-mediated decay (NMD) pathway is another quality control checkpoint for aberrant transcripts in the cytoplasm, and recently emerged as a key player for fine-tuning of both coding and noncoding gene expression (Smith and Baker 2015). A genome-wide survey of human lncRNA sequences showed that most lncRNAs harbor short open reading frames (ORFs) that would lead to activation of the NMD pathway (Niazi and Valadkhan 2012). In fact, subsets of lncRNAs found in budding yeast, the model plant *Arabidopsis* and animals are subject to degradation through the NMD pathway (Kurihara et al. 2009; Tani et al. 2013; Ruiz-Orera et al. 2014).

The cytoplasmic exonuclease Xrn1 is the final enzyme responsible for the degradation of de-capped and de-adenylated transcripts that have been recognized and processed by NMD components. The deletion of *XRN1* in budding yeast also leads to the accumulation of more than a thousand cryptic noncoding transcripts termed ‘XUTs’ (<u>X</u>rn1-sensitive <u>u</u>nstable <u>t</u>ranscripts), most of which are distinct from the noncoding transcripts that arise by exosome depletion (van Dijk et al. 2011). Many XUTs are antisense to annotated genes and seemed to have repressive roles in sense transcription by modulating chromatin status of the promoter regions (van Dijk et al. 2011).

It has been argued that organismal complexity is correlated with expression dynamics of noncoding transcripts (Mattick et al. 2010; Liu et al. 2013; Gaiti et al. 2015; Quinn and Chang 2015). In the Kingdom Fungi, multicellular fruiting bodies have independently evolved at least twice in the diverging lineages (Knoll 2011; Nguyen et al. 2017). Given the key regulatory roles of lncRNAs in sexual sporulation and morphological transition in yeasts (Bumgarner et al. 2009; Chacko et al. 2015), lncRNAs may have exerted their roles in evolution of multicellularity and sexual development in filamentous fungi. *Fusarium graminearum* is a plant pathogenic fungus infecting our staple crops, such as wheat and corn, and thus has been a model for studying the developmental process of perithecia, the sexual fruiting bodies of the fungus, as well as other interesting aspects of biology including host-pathogen interaction and mycotoxin production (Trail 2009; Ma et al. 2013). The fungus is probably the best organism for investigating the lncRNA catalog in fruiting body-forming fungi, as perithecia develop at sufficient synchronicity in culture media, enabling time-series transcriptome analyses with this microscopic organism (Hallen et al. 2007; Sikhakolli et al. 2012; Trail et al. 2017). Also, the genome sequence assembly is complete, featuring a total 4 chromosomes (Cuomo et al. 2007), and the genome has been annotated and curated—although it still lacks lncRNA annotations (King et al. 2015, 2017). In addition, plentiful genetic resources have accumulated through large-scale functional studies of perithecia development (Son et al. 2011; Wang et al. 2011a; Yun et al. 2015; Kim et al. 2015b; Liu et al. 2016; Son et al. 2017; Trail et al. 2017).

The goal of the present study was to characterize lncRNAs that are specifically expressed in the fungal fruiting body undergoing sexual development and to investigate their developmental stage-specific expression and their regulation. We identified lncRNAs with a pipeline that constructs *de novo* transcript annotations by combining RNA-seq data from vegetative and sexual developmental stages, and then removes those with detectable protein-coding potential or monotonous expression profile. Hundreds of lncRNAs that exhibit dynamic expression patterns were found, thereby expanding the universe of genomes known to have significant noncoding roles in development—specifically, here, the multicellular development of fungal fruiting bodies.

## RESULTS

### Transcriptional reprogramming during perithecia formation

To obtain time-course transcriptome data during the sexual development of *F. graminearum*, we sequenced samples from hyphae, strands of cells that make up vegetative stages of most of fungi (S0), and from five successive sexual stages (S1-S5; Fig. 1A) that capture key morphological transitions during the perithecia development, defined at the formation of: S1—formation of perithecial initials (hyphae curls that give rise to the perithecial tissues), S2—perithecial walls, S3—paraphyses (sterile cells supporting perithecia), S4—asci (sac-like structures in which ascospores develop), and S5—ascospore maturation (Trail and Common 2000). A total of 480 million RNA-seq reads were generated from 18 samples (6 stages × 3 replicates), and there were average 25 million mapped reads per sample (Supplemental Fig. S1). We validated our sampling scheme by perithecial morphology for 3 biological replicates, using the BLIND program (Anavy et al. 2014) that determined the sequence of the developmental time-course data without prior information other than gene expression data (Supplemental Fig. S2).

Differentially-expressed (DE) genes between any two successive developmental stages (>4-fold at 5% false discovery rate [FDR]) were mostly unique when compared to other pairwise stage comparisons (Fig. 1B). Overrepresented GO terms for the stage-specific DE genes reflected key biological processes during the morphological transitions (Supplemental Table S1). For example, the GO term ‘lipid metabolism’ had the highest representation in the ‘S0 vs. S1’ comparison, although statistically non-significant (Fig. 1C). The accumulated lipids in hyphae and perithecial initials are vital for paraphyses and asci development (Guenther et al. 2009). Perithecia dramatically increased in size and became more rigid during S2 and S3, which is accompanied by GO terms related to carbohydrate metabolisms (Fig. 1C). Finally, when the asci develop from the fertile layer during S3 and S4, meiosis-related genes were significantly enriched for GO terms including ‘meiotic cell cycle process’ and ‘ascospore formation’ at 5% FDR (Fig. 1C). It is noteworthy that most of the DE genes were upregulated in the later developmental stages (Supplemental Fig. S3), indicating that gene activation is the most common means of gene regulation during the sexual development.

**Figure 1.**
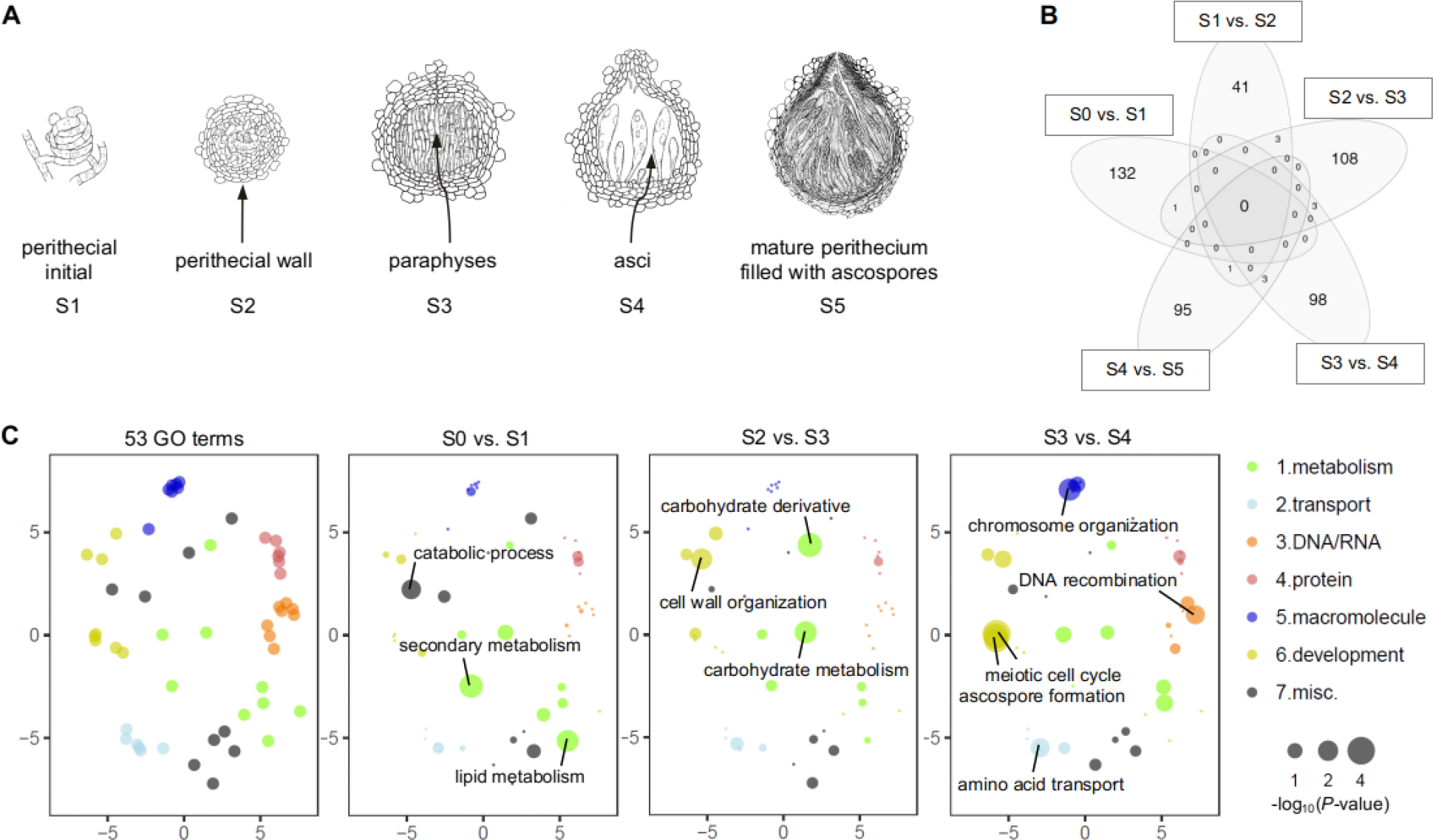
Transcriptome of the *F. graminearum* perithecia. (*A*) Emergence of new tissues at the defined developmental stages during the perithecia formation (S1–S5; not drawn to scale). (*B*) Venn diagram showing the number of differentially expressed (DE) genes between two successive developmental stages (>4-fold; 5% FDR). Note that most of the DE genes were unique in each comparison. (*C*) Functional enrichment analyses for DE genes between two successive developmental stages. Fifty-three GO terms— that can be broadly categorized into 7 biological processes—were assessed for degree of functional enrichment (Supplemental Table S1), and were projected to two-dimensional semantic spaces. Only GO terms with *P* < 0.05 were depicted in each panel.

### Identification of lncRNA in perithecia

To discover lncRNAs expressed during the perithecia development, we adopted an established protocol for novel transcripts identification, with some modifications (Weirick et al. 2016; Supplemental Fig. S4). First, we constructed *de novo* transcript annotations (28,872 transcripts expressed from 20,459 genomic loci), and identified potentially novel transcripts that were absent in the reference annotations (Table 1). For noncoding transcripts identification, coding potential of the novel transcripts was computed by using the CPAT program (Wang et al. 2013). To maximize both sensitivity and specificity for noncoding transcript detection, the program was trained on the *F. graminearum* genome dataset and the threshold was set to a CPAT score of 0.540 (Supplemental Fig. S4) (*cf*. 0.364 for humans, 0.440 for mice; Chakraborty et al. 2014). The transcripts with a low coding potential (CPAT score ≤ 0.540) were further scanned against Pfam and Rfam databases to filter out transcripts encoding protein domain(s) and harboring any known structural RNA motifs, respectively (*E* < 10^−10^; Table 1). Finally we only retained transcripts that were differentially expressed in at least one developmental stage (5% FDR), yielding a total 547 lncRNA candidates (Table 1; Supplemental Table S2).

**Table 1.** Identification of lncRNAs expressed during sexual development

| Transcript class (code)<sup>a</sup> | Novel transcripts<sup>b</sup> | Noncoding transcripts<sup>c</sup> | Differentially expressed transcripts<sup>d</sup> |
| --- | --- | --- | --- |
| Splicing variants ('J') | 5,185 | 92 | 17 |
| Tandem transcripts ('P') | 740 | 388 | 100 |
| Intergenic transcripts ('U') | 2,215 | 1,054 | 167 |
| Antisense transcripts ('X') | 2,892 | 1,040 | 263 |
| Sum | 11,032 | 2,574 | 547 |

<sup>a</sup> Transcript class codes were tagged by the *gffcompare* program (Pertea et al. 2016).
<sup>b</sup> Among total 11 transcript class codes, transcripts tagged with the class codes 'J', 'P', 'U', and 'X' were considered as novel transcripts (see Supplemental Methods).
<sup>c</sup> Noncoding transcripts were identified by the coding potential assessment tool program (Wang et al. 2013), and further filtered by Pfam and Rfam database searches.
<sup>d</sup> Differentially expressed noncoding transcripts at least one developmental stage were identified as *F. graminearum* lncRNAs.

The identified lncRNAs were distributed across the 4 chromosomes, and generally shorter with fewer exons, when compared to mRNAs (transcripts with CPAT score > 0.540) (Fig. 2A-C). Based on the relative position to mRNAs, lncRNAs can be classified as antisense lncRNA (ancRNA) or long intergenic ncRNA (lincRNA). There were 280 ancRNAs that overlapped more than 100 bp of an mRNA on the opposite strand, and 237 lincRNAs that were situated between annotated genes (Supplemental Table S2). The mean AU content of ancRNA and lincRNA sequences falls between the coding sequences of the mRNAs and the intergenic regions (Fig. 2D). These distinctive genomic features of the lncRNA sequences are commonly observed in other eukaryotes (Nam and Bartel 2012; Pauli et al. 2012; Gaiti et al. 2015; Li et al. 2016; Nyberg and Machado 2016). However, we identified only 5 lncRNAs that showed similarity with lncRNAs in other eukaryotes (*E* < 10^−10^; Supplemental Table S3), suggesting either the poor status of lncRNA annotations in filamentous fungi or a high degree of sequence divergence in fungal lncRNAs.

**Figure 2.**
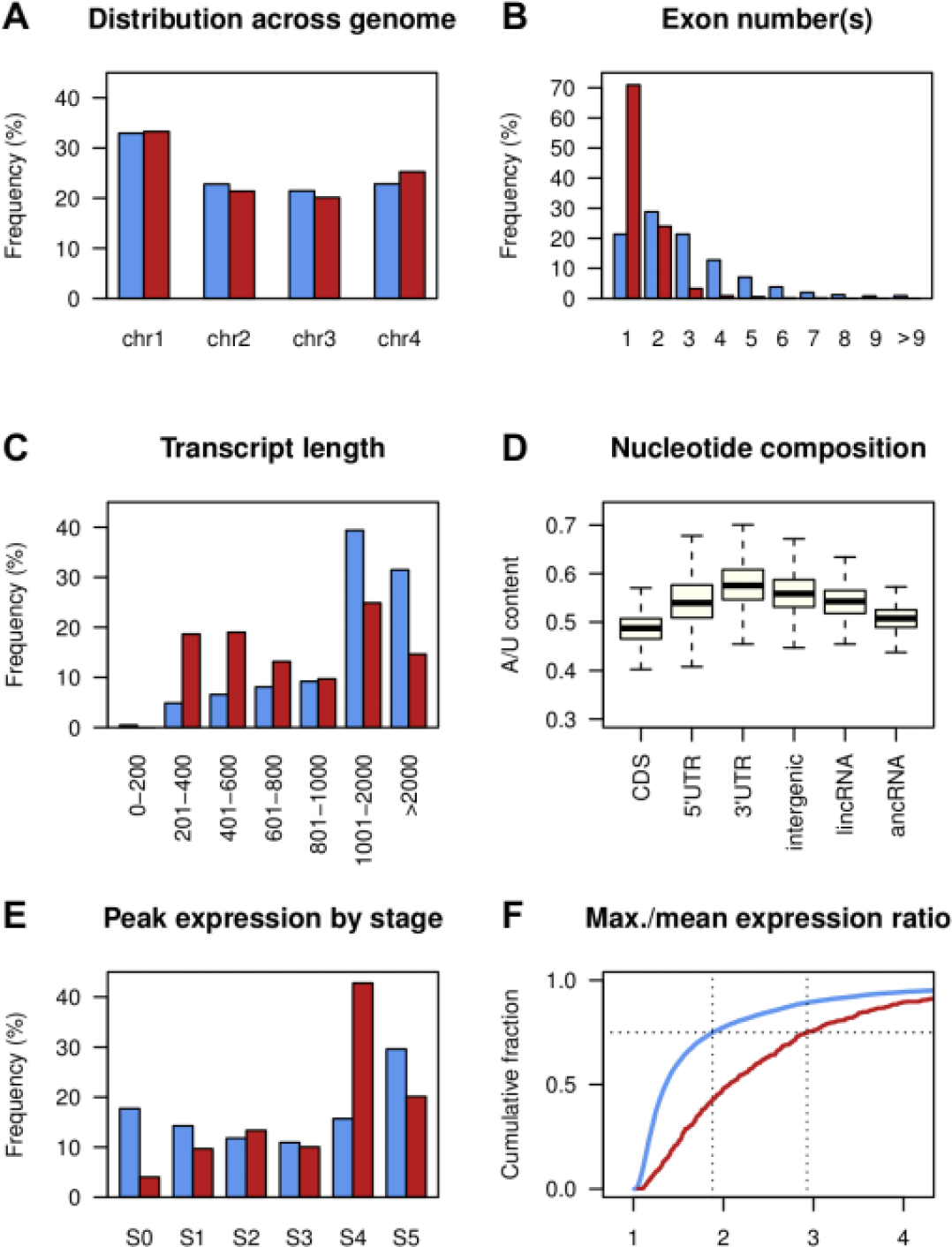
Genomic features of *F. graminearum* lncRNA. (*A*) Distribution of mRNA and lncRNA across the four chromosomes. (*B*) Distribution of exon numbers per transcript. (*C*) Transcript length distribution. (*D*) A/U content of mRNA coding regions (CDS), 5′ UTRs, 3′ UTRs, intergenic regions, long intergenic lncRNA (lincRNA), and antisense lncRNA (ancRNA). Box and whisker plots indicate the median, interquartile range between 25th and 75th percentiles (box), and 1.5 interquartile range (whisker). (*E*) Distribution of developmental stages at which mRNA and lncRNA showed the highest expression level. (*F*) Cumulative distributions of ratios of maximum and mean expression values across the developmental stages. mRNA—blue boxes or line, and lncRNA—red boxes or line.

### Developmental expression of lncRNA

The sexual stage transcriptome data showed predominance of lncRNAs at the meiotic stage (S4) where the expression of many lncRNAs peaked (234 out of the 547 lncRNAs; Fig. 2E). We compared the degree of differential expression of lncRNAs to that of mRNAs (9,457 transcripts differentially expressed in at least one developmental stage at 5% FDR). The ratio of the maximum expression among six developmental stages to the mean expression over the remaining five stages was calculated for lncRNAs and the differentially expressed mRNAs. By this metric, lncRNA was prone to be more differentially expressed than mRNA (*P* = 2.2 × 10^−16^, KS-test statistic *D* = 0.39), with the third quartile value of the ratio measuring 2.93 for lncRNA and 1.88 for mRNA (Fig. 2F). Also, we identified seven co-expressed clusters of lncRNAs that showed developmental-specific expression patterns, suggesting distinct roles of lncRNAs in different stages of perithecia development (Fig. 3A).

**Figure 3.**
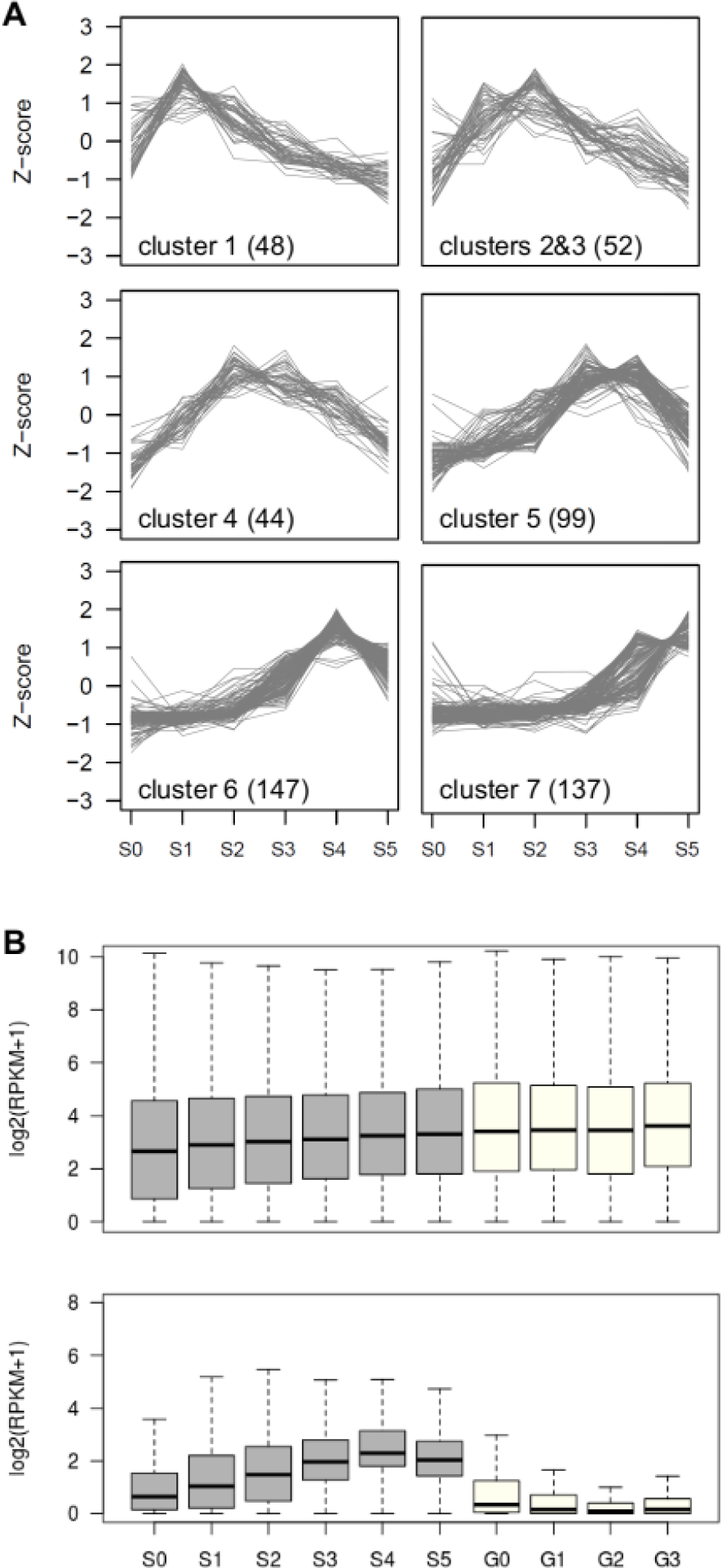
Sexual stage-induced lncRNA in *F. graminearum*. (*A*) Co-expressed clusters of lncRNAs. Trend plots of Z-score normalized expression values for lncRNAs (numbers in parenthesis) in a given cluster were presented. (*B*) Expression distribution of mRNA (upper panel) and lncRNA (lower panel) for the sexual development transcriptome (grey boxes) and the vegetative growth transcriptome (white boxes). Box and whisker plots indicate the median, interquartile range between 25th and 75th percentiles (box), and 1.5 interquartile range (whisker).

In addition to the sexual stage dataset, we obtained transcriptome data during spore germination to investigate the degree of lncRNA expression in vegetative stages. The dataset was comprised of four spore germination stages: G0—fresh spore, G1—polar growth, G2—doubling of long axis, and G3— branching of hyphae (Supplemental Fig. S5). Overall expression of the lncRNA gradually increased over the course of perithecia development, peaking at S4, while most of the lncRNA remained unexpressed or had low expression in the germination stages, indicating that most of the lncRNA expression is sexual stage-specific (Fig. 3B).

### Verification of lncRNA production

To validate lncRNA expression, we chose eight lncRNAs and performed 3’ RACE-PCR and Sanger sequencing. All the selected lncRNAs were amplified from total RNA extracts and their polyadenylation sites were determined (Supplemental Fig. S6). Also, to examine if there is an intraspecific conservation in the lncRNA content and expression, we utilized degradome-seq data of another *F. graminearum* wild-type strain sampled at meiotic stage (Son et al. 2017). Degradation of lncRNA transcripts expressed at S4 was evident, which in turn confirmed the consistent lncRNA production in the two strains (Fig. 4; also see below, Fig. 5B).

**Figure 4.**
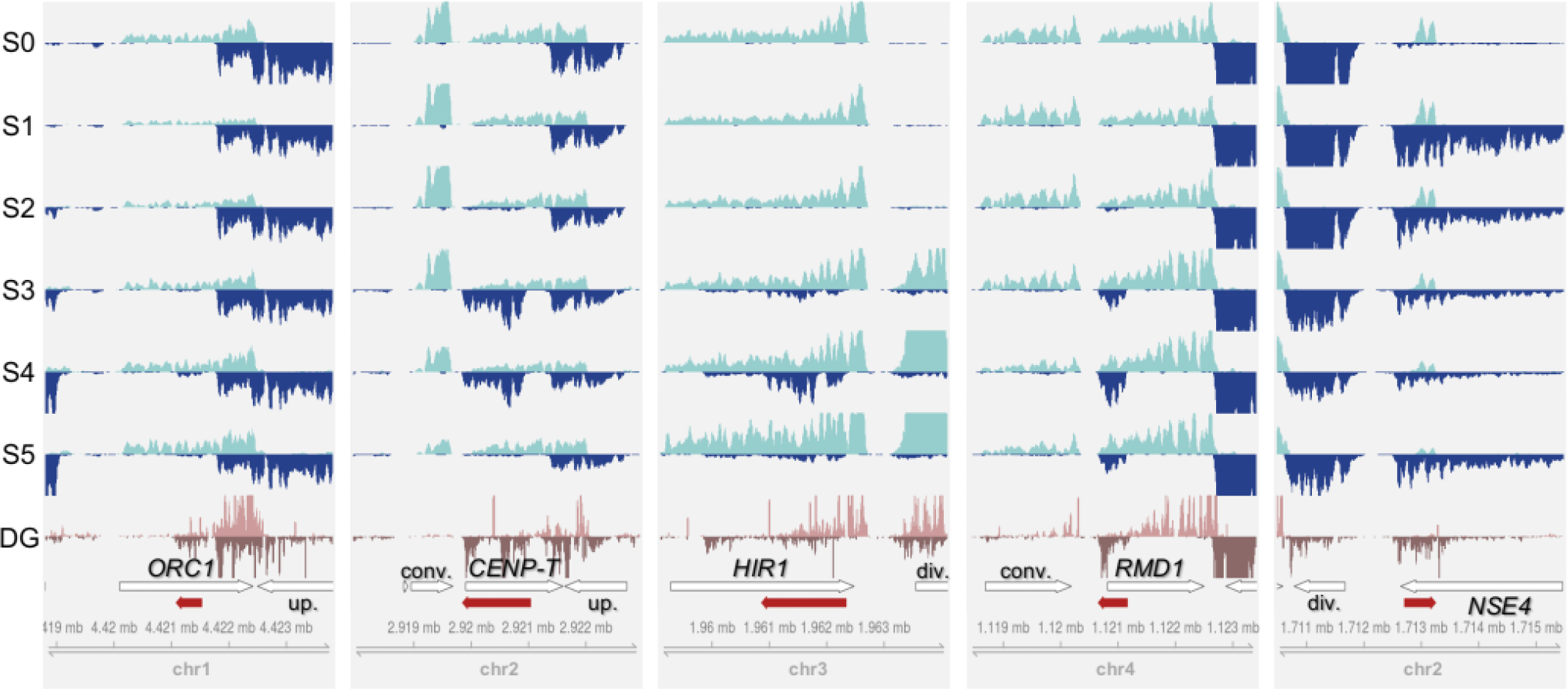
Examples of lncRNA expression across the sexual development. Per-base coverage of transcripts was plotted for both DNA strands in a 5 kb window. For the perithecia transcriptome datasets (S0–S5), mapped reads of 3 biological replicates were pooled, then subsampled to 60 million reads for visual comparison of expression levels across the stages. For the degradome-seq datasets (DG), mapped reads of 2 replicates were pooled and displayed. The positions of lncRNAs (red arrows) and their neighboring genes (white arrows) are shown in the annotation track with genome coordinate at the bottom of each panel. The genes overlapping lncRNAs on the opposite strand are labeled with abbreviated gene names in bold: *CENP-T*—<u>cen</u>tromere <u>p</u>rotein <u>T</u> (FGRRES_16954), *HIR1*—<u>hi</u>stone regulatory protein 1 (FGRRES_05344), *NSE4*—<u>n</u>on-<u>s</u>tructural maintenance of chromosome <u>e</u>lement 4 (FGRRES_17018), *ORC1*—<u>o</u>rigin recognition <u>c</u>omplex subunit 1 (FGRRES_01336), *RMD1*—required for <u>m</u>eiotic <u>d</u>ivision 1 (FGRRES_06759). In relation to lncRNA position, neighboring genes are also labeled as follows: div.— divergently transcribed gene on the opposite strand, conv.—convergently transcribed gene on the opposite strand, up.—upstream gene in tandem on the same strand.

**Figure 5.**
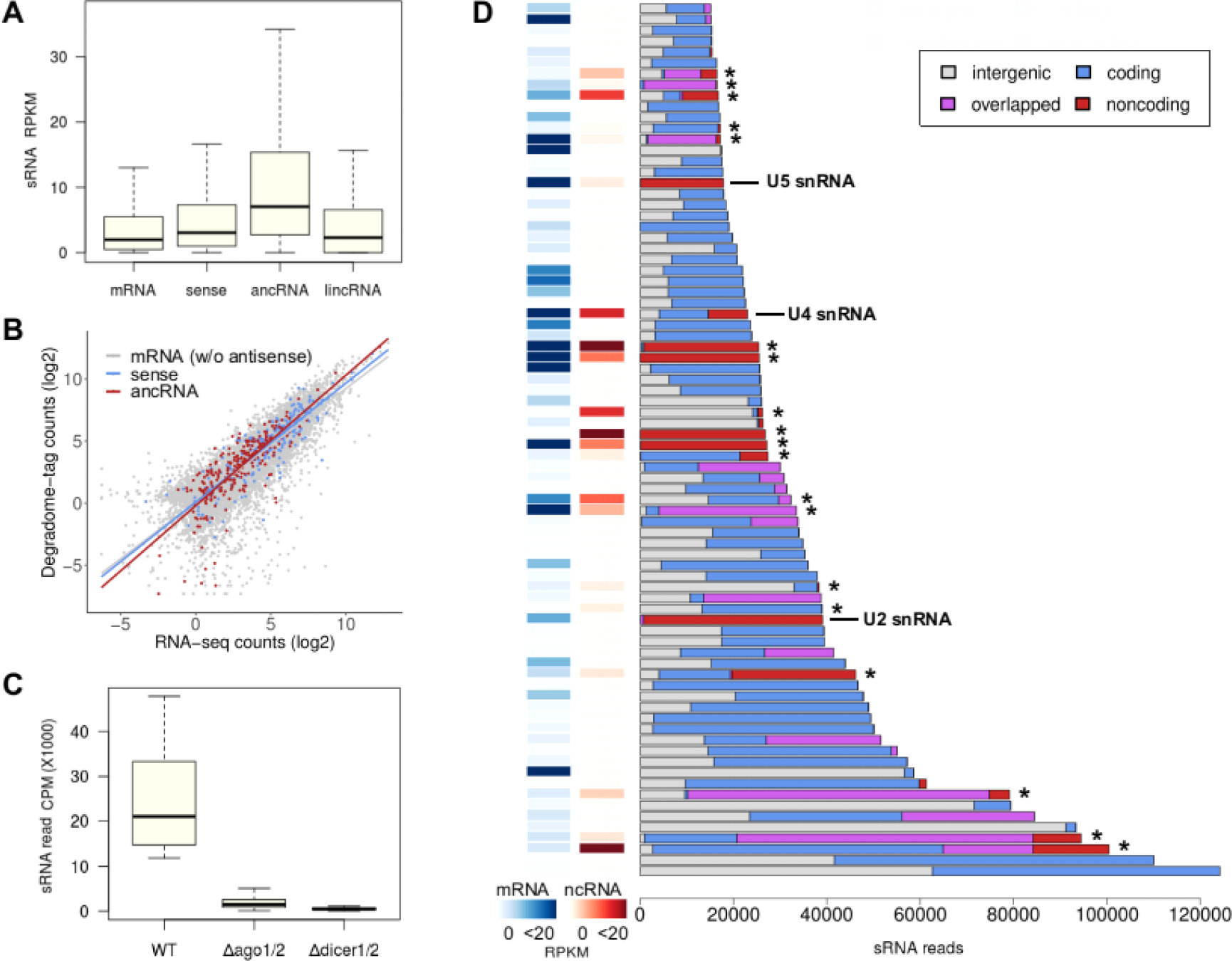
lncRNA associated with small RNA-enriched loci. (*A*) sRNA reads mapped to mRNAs without antisense transcripts (10,928 loci), sense mRNAs for ancRNAs (295), ancRNAs (276), and lincRNA (235) were represented as RPKM. The number of sRNA reads aligned to ancRNA loci was more than the other classes of transcripts (Benjamini-Hochberg adjusted *P* < 0.0001, Dunn’s pairwise multiple comparisons). (*B*) Correlation analysis between transcript abundance and degradome-tag count at meiotic stage. Lines depict regressions for different classes of transcripts. (*C*) sRNA reads mapped to top 80 sRNA clusters in different genotypes were represented as counts-per-million (CPM). The number of sRNA reads mapped to the clusters were significantly reduced in Δ*dicer1/2*, the double-deletion mutants of Dicer genes (FGRRES_09025 and FGRRES_04408), and Δ*ago1/2*, the double-deletion mutants of Argonaute genes (FGRRES_16976 and FGRRES_00348) (Son et al., 2017). (*D*) Fractions of sRNA reads mapped to intergenic regions (white), coding genes (blue), overlapped regions (purple), and noncoding genes (red) including lncRNAs marked with asterisks in top 80 sRNA clusters. Overlapped regions were defined if coding or noncoding genes in the region were present on the both sides of DNA in the *de novo* annotations. Expression values (RPKM) for the closest coding gene (mRNA) to the center of each sRNA cluster are shown as heat maps, along with expression values for ncRNAs, if any, that were present in the same cluster.

Fungal genomes are known for having shorter intergenic spaces, compared to other eukaryotic genomes. Therefore, fungal lncRNAs could be a transcriptional noise arising from neighboring genes. To test this, we examined global patterns of the expression correlation between lncRNA and neighboring genes, with close examination of some selected examples whose expression was confirmed by 3’ RACE-PCR. The lncRNAs antisense to *ORC1*, *ORC2*, and *CENP-T* each had an upstream gene in close proximity on the same strand (Fig. 4; Supplemental Fig. S6). However, the sexual stage expression between the lncRNAs and their respective upstream genes was not correlated (| *r* | < 0.50; Supplemental Table S4). Positive correlation was observed in expression levels between lncRNAs and their divergently transcribed genes as in the *HIR1* and *NSE4* loci (*r* > 0.8; Fig. 4; Supplemental Table S4), indicating prevalence of bidirectional promoters for lncRNA transcription (Neil et al. 2009; Xu et al. 2009; Pelechano et al. 2013). On the other hand, the expression of lncRNAs and their convergently transcribed genes tend not to be correlated, as in the *CENP-T* and *RMD1* loci (| *r* | < 0.50; Fig. 4; Supplemental Table S4). These patterns were globally observed in lncRNA-associated loci (Supplemental Fig. S6), suggesting that the lncRNAs were not likely to be misannotated extensions of neighboring genes (Cabili et al. 2011; Ulitsky et al. 2011; Nam and Bartel 2012).

### Identification of sRNA-enriched loci associated with lncRNAs

We found 300 sense mRNA–ancRNA pairs with different orientations: 5’-to-5’ partial (70), 3’- to-3’ partial (61), and full overlaps (169). One of the most common mechanisms involving antisense transcription is the RNAi pathway incorporating sRNAs generated from the double-stranded RNA regions. To investigate the degree and effect of sRNA production in the ancRNA loci, we analyzed the previously published sRNA-seq and degradome-seq data at meiotic stage (Son et al. 2017). As expected, sRNA reads were mapped at a higher frequency to ancRNAs than to mRNAs without overlapping antisense transcripts, the sense mRNAs or lincRNAs (Fig. 5A; *P* < 1.2 × 10^−8^, Kruskal-Wallis test statistic *H* = 155), suggesting that the ancRNA loci may serve as a major source for endogenous sRNA production. However, the correlation of the degradome-seq and our RNA-seq data at S4 showed that sRNA-mediated endonucleolytic cleavage of ancRNAs and sense mRNAs was comparable to that of mRNAs without antisense transcripts (Fig. 5B), implying that the ancRNA loci were not preferentially targeted by RNAi machinery, post-transcriptionally.

It remains paradoxical that gene silencing induced by heterochromatin formation requires sRNA production via co-transcriptional processes, sometimes from lncRNAs (Motamedi et al. 2004; Bühler et al. 2006; Zhang et al. 2008; Bayne et al. 2010; Dang et al. 2016). To search for any lncRNAs associated with sRNA-enriched loci that could be indicative of transcriptional gene silencing events, we examined top 80 sRNA clusters ranked by the number of mapped reads, which accounted for 62% of mapped sRNA reads (Supplemental Table S5). Production of sRNAs in the top 80 clusters was dependent on Dicers and Argonauts, indicating that the sRNAs were produced by RNAi machinery (Fig. 5C). Most of the sRNA clusters were found in genic regions, containing at least one annotated gene, to which a large portion of sRNA reads were mapped (Fig. 5D). We observed that the coding genes closest to the centers of sRNA clusters exhibited overall low expression (RPKM < 0.5 in 22 out of the 80 clusters) (Fig. 5D). A significant portion of sRNAs were also derived from noncoding transcripts and genic regions overlapped with antisense transcripts, some of which were identified as lncRNAs (Fig. 5D; Supplemental Fig. S7). Unexpectedly, the lncRNAs associated with sRNA clusters exhibited moderate expression (*n* = 19, median expression 5.2 in RPKM). In addition, the coding genes closest to the centers of the sRNA clusters showed higher expression levels (*n* = 19, median expression 4.0), compared to those without an associated lncRNA (*n* = 61, median expression 1.3; *P* = 0.029, Mann-Whitney test statistic *U* = 747).

### Co-expression of lncRNAs and their sense transcripts

We could not find strong evidence for sRNA-mediated transcriptional and posttranscriptional gene silencing in the ancRNA loci. Interestingly, we did observe gene expression correlation in many ancRNAs and sense mRNA pairs across the sexual stages (Fig. 6; 85 out of the 300 pairs with Pearson’s correlation | *r* | > 0.70; Fisher’s exact test, *P* < 0.05), most of which were positively correlated (76 out of the 85 pairs). We asked whether the ancRNAs are antisense to genes involved in a specific biological process. Notably, the positively correlated sense mRNAs were enriched for the GO term ‘DNA metabolism’ at 5% FDR (Supplemental Table S6). This observation might be a consequence of the underlying structure of the dataset, in which the GO term was also the most overrepresented GO term for all the sense mRNAs overlapping the 280 lncRNAs (although this observation was not statistically significant, applying a 5% FDR; *P* = 0.006).

**Figure 6.**
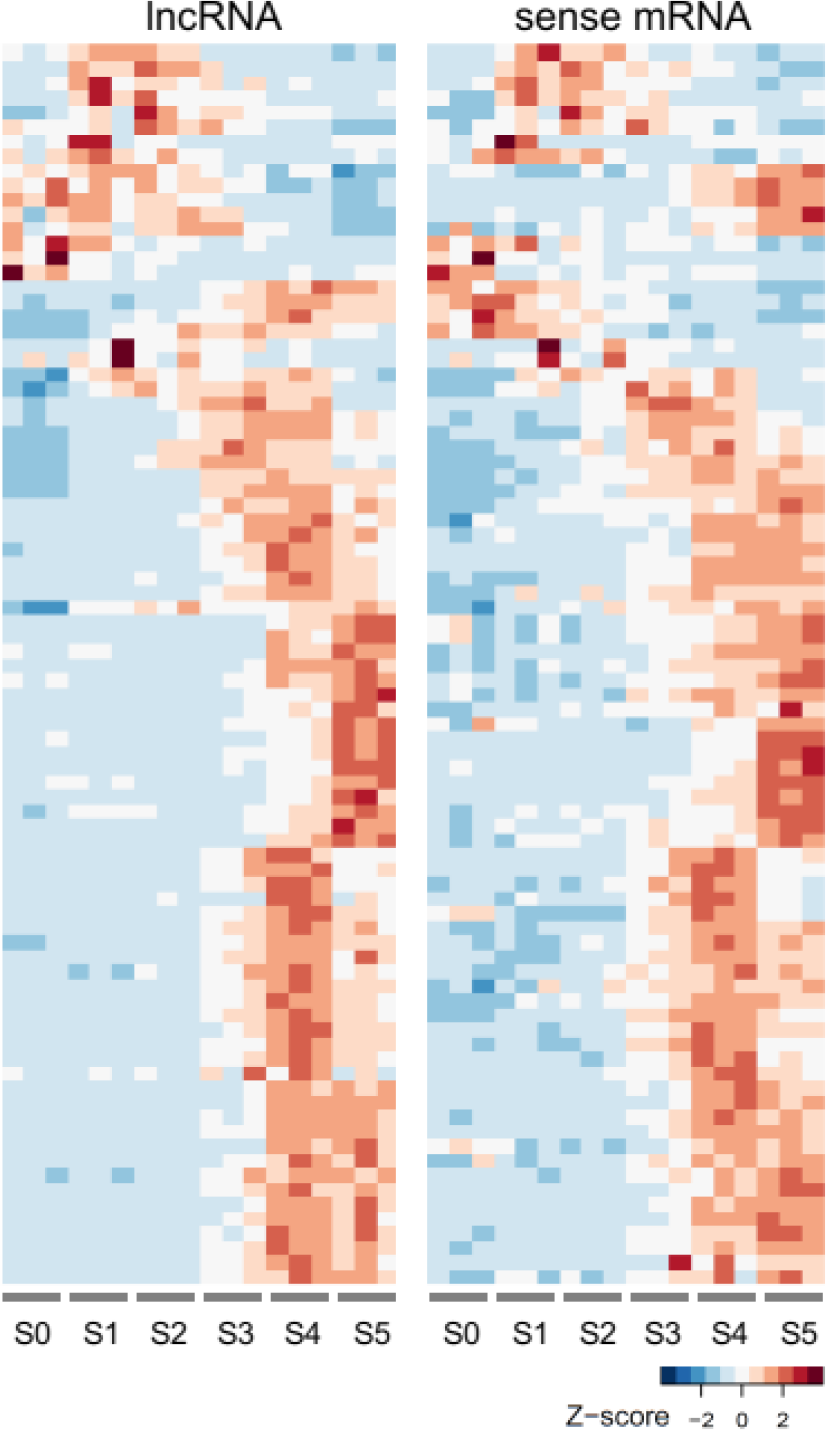
Parallel induction of sense mRNA and antisense lncRNA pairs during sexual development. Expression data of sense mRNA and antisense lncRNA pairs in 18 samples for the perithecia transcriptome (S0–S5) were re-ordered by the BLIND program (Supplemental Fig. S2). The RPKM values of sense mRNA and antisense lncRNA pairs with absolute Pearson’s correlation greater than 0.7 (*P* < 0.05; Fisher’s exact test) were clustered by Euclidean distance and heat maps of Z-score normalized RPKM values were presented.

### Regulation of sexual development and lncRNA expression by NMD

The exosome and NMD components regulate the expression of noncoding genes as well as coding genes for sexual sporulation in yeasts. To better understand how lncRNA expression was regulated during the perithecia development, we initially aimed to dissect the roles of two exonuclease genes (*RRP6*: FGRRES_06049 and *XRN1*: FGRRES_06799), and two RNA helicase genes functionally associated with both the exosome and NMD pathway (*DBP2*: FGRRES_16145 and *MTR4*: FGRRES_01656) (Cloutier et al. 2012; Wery et al. 2016). However, efforts to obtain deletion strains lacking either the *DBP2*, *MTR4* or *RRP6* were unsuccessful (Supplemental Fig. S8), presumably due to the synthetic lethality of the genes when deleted, as with *Neurospora crassa* (Cheng et al. 2005; Emerson et al. 2015). However, we were able to generate Δ*xrn1* strain in *F. graminearum* and observed delayed perithecia development and deformed ascospores with variable sizes (Fig. 7A; Supplemental Fig. S8). Similarly, defects in ascospore formation were observed in heterokaryotic Δ*dbp2* strain containing wild-type nuclei (Fig. 7A), suggesting that maintaining RNA homeostasis by NMD pathway is crucial for meiotic divisions and ascospore formation.

**Figure 7.**
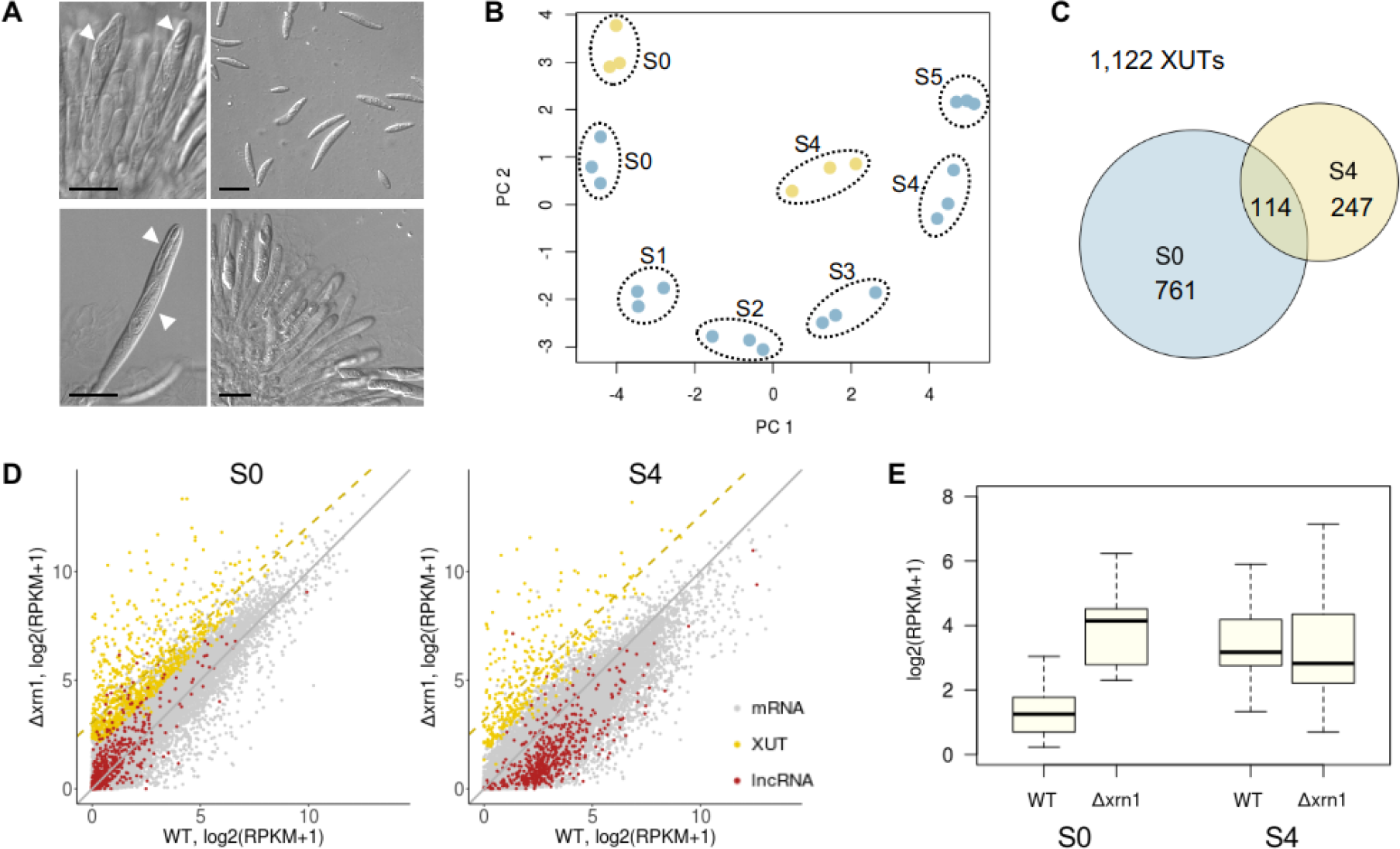
Sexual stage-induced lncRNAs regulated by the NMD pathway. (*A*) Asci and ascospores of Δ*xrn1* (upper panels) at 11 days after sexual induction and heterokaryotic Δ*dbp2* strain (lower panels) at 9 days after sexual induction. Arrows indicate uneven delimitation of the ascospore membrane in the asci, resulting in deformed ascospore production (right panels). Scale bar = 20 μm. (*B*) Principal components analysis on the perithecia transcriptome data of the wild-type (S0–S5, blue circles) and Δ*xrn1* transcriptome data (S0 and S4, yellow circles). (*C*) Venn diagram showing the overlap of <u>X</u>rn1-sensitive <u>u</u>nstable <u>t</u>ranscripts (XUTs) at S0 and S4. (*D*) 2D-plots of different classes of transcripts between the wild-type (abscissa) and Δ*xrn1* (ordinate) expression data at S0 and S4 (diagonal—grey line; regression for the XUTs—yellow dashed line). (*E*) Expression distribution of 25 lncRNAs that were identified as XUTs at S0. Box and whisker plots indicate the median, interquartile range between 25th and 75th percentiles (box), and 1.5 interquartile range (whisker).

To identify lncRNA whose expression was controlled by NMD activity, we analyzed the transcriptome data from the wild-type (WT) and the Δ*xrn1* strain at hyphae stage (S0) and meiotic stage (S4). The transcriptome data of the Δ*xrn1* strain were distinct, but most similar, to that of WT at the corresponding stages, indicating drastic effects of *XRN1* on gene expression levels as a major component for RNA turnover (Fig. 7B). After expression values were normalized with 124 ribosomal protein genes that were known to be relatively insensitive to NMD activity (van Dijk et al. 2011), we identified a total of 1,122 XUTs that were differentially expressed at S0 or S4 in Δ*xrn1* (>3-fold increase at 5% FDR; Fig. 7C; Supplemental Table S7). After the normalization, the XUTs identified at S0 and S4 showed a median 5.0- and 5.8-fold increase in Δ*xrn1*, respectively (Fig. 7D; Supplemental Table S7). Many XUTs were previously annotated transcripts or isoforms of them (70%; 781/1,122), and only 11% of XUTs (122/1,122) were predicted to be noncoding transcripts (CPAT score ≤ 0.540) (Supplemental Table S7). Among the noncoding XUTs, we identified 25 lncRNAs whose expression was elevated upon the Xrn1 depletion at S0 and showed increasing patterns across the sexual stages (Fig. 7E; Supplemental Fig. S9). In addition, many of the coding XUTs identified at S0 were also induced as the sexual development progressed (Supplemental Fig. S10). Interestingly, we found that key components of RNA-induced silencing complex (RISC; Chen et al. 2015) were identified as XUTs, such as *DICER2* (FGRRES_04408), *AGO1* (FGRRES_16976), *N. crassa QIP* homolog (FGRRES_06722), and *RDRPs* (RNA-dependent RNA polymerases: FGRRES_01582, 04619, and 09076) (Supplemental Table S7).

## DISCUSSION

Here we profiled transcriptomes of vegetative and sexual stages that span the entire life cycle of *F. graminearum*. lncRNAs are usually an order of magnitude less abundant than mRNAs (Mercer et al. 2011; Cabili et al. 2015), so the unprecedented depth of the sequencing data we generated (total 938 million mapped reads) enabled us to capture scantly expressed noncoding transcripts. This study has revealed global properties of lncRNAs during the perithecia development, characterized by dynamic and developmental stage-specific expression. In the last step of our lncRNA identification pipeline, we only included differentially expressed noncoding transcripts to discern lncRNAs of biological significance. This filtering step allowed us to identify low-abundance lncRNAs that alone could be argued to be transcriptional noise (ENCODE Project Consortium 2007; Bakel et al. 2010). For many antisense lncRNAs, we detected parallel induction with their respective sense transcripts across the sexual development and identified a subset of lncRNAs that are sensitive to nonsense-mediated decay activity before sexual induction.

Our lists of *F. graminearum* lncRNAs contain many confident lncRNA annotations, but are still far from complete. It is primarily due to difficulty in unequivocal determination of whether a transcript is coding or noncoding (Anderson et al. 2015). Although protein products from most of the bonafide lncRNAs with predicted ORFs have not been detected in cells (Bánfai et al. 2012; Gascoigne et al. 2012), many lncRNAs with one or more ORFs have been shown to be associated with ribosomes in budding yeast, *Arabidopsis*, and animals (Wilson and Masel 2011; Chew et al. 2013; Ingolia et al. 2014; Juntawong et al. 2014; Ji et al. 2015; Carlevaro-Fita et al. 2016). By scrutinizing the sRNA-enriched loci, we identified as lncRNAs several transcripts with short ORFs (ranging from 49 to 227 amino acids). These novel transcripts were initially filtered out by the CPAT program, however another annotation tool, CPC2 (Kang et al. 2017) classified them as lncRNAs, and the lncRNA annotations were also supported by the lack of cross-species conservation of the deduced polypeptides (*E* ≥ 10^−10^; Supplemental Table S8).

Another source of false-negatives in our dataset may have arisen from poly-A-based library preparation that excluded some lncRNAs lacking poly-A tail. This may account for the inclusion of fewer noncoding transcripts in our XUT identification process. More precise identification of lncRNAs, especially for those undergoing deadenylation by the RNA quality control machineries, should be accompanied by cap-analysis gene expression (CAGE)-seq (Bogu et al. 2016; Wery et al. 2016; Liu et al. 2017). However, most lncRNAs identified in this study were also detected in the degradome-seq (Son et al. 2017), another sequencing method (*aka*. parallel analysis of RNA ends; German et al. 2008), which further validates their authenticity.

To better understand how the expression of lncRNAs was controlled during perithecia development, we identified XUTs, as the NMD pathway has been shown to determine the fate of lncRNAs (Smith and Baker 2015; Wery et al. 2016). However, only 5% of the sexual stage-induced lncRNAs (25/547) were affected by Xrn1 activity, and the majority of the lncRNAs seemed to escape from the RNA surveillance system. The proportion of XUTs among the lncRNAs in *F. graminearum* is comparable to that in other eukaryotes, where approximately 4–14% of noncoding transcripts are known to undergo NMD (Ruiz-Orera et al. 2014). Interestingly, several sexually-induced RISC components were highly upregulated in Δ*xrn1*, indicating a possible link between the gene silencing pathway and the NMD pathway. Further studies on lncRNAs and the RNAi machinery that were affected by NMD activity will provide novel insights into regulatory mechanisms via altered RNA metabolism during the sexual development.

We were unable to test the role of the exosome components in lncRNA expression, as the major exosome component (*RRP6*) and the functionally-related helicase genes (*DBP2* and *MTR4*) all seemed to be essential in *F. graminearum*. In fission yeast, the nuclear exosome mediates gene silencing of coding and noncoding loci during vegetative growth, and the sexual sporulation is triggered by disassembly of heterochromatin on the silenced loci (Zofall et al. 2012). Therefore, it will be interesting to see if the regulation of lncRNA expression is achieved by modulation of chromatin status before and after the sexual induction in *F. graminearum*.

Antisense transcription can influence synthesis, expression kinetics, and stability of sense transcripts through a variety of mechanisms (Böhmdorfer and Wierzbicki 2015; Quinn and Chang 2015). Antagonism of lncRNAs against meiotic gene expression has been documented in yeasts (Hongay et al. 2006; Ni et al. 2010; Rhind et al. 2011; Zhang et al. 2011; Chen et al. 2012; van Werven et al. 2012). Nevertheless, there has been growing evidence that lncRNAs activate repressed genes by modulating local chromatin structures, which can facilitate coordinated gene expression in budding yeast (Cloutier et al. 2013, 2016) and in other eukaryotes (Ørom et al. 2010; Wang et al. 2011b; Li et al. 2013; Boque-Sastre et al. 2015). In favor of this regulatory phenomenon, we hypothesize that the *F. graminearum* lncRNAs, which showed parallel induction with sense transcripts across the sexual development, may play a role in regulation of DNA synthesis and degradation at later developmental stages, according to “guilt-by-association” (Guttman et al. 2009; Gaiti et al. 2015). However, lncRNAs often exhibited cell type-specific expression (Bitton et al. 2011; Bumgarner et al. 2012; Li et al. 2013), and even allele-specific expression within the same nucleus (Rosa et al. 2016). Therefore, we cannot rule out the possibility that the positively correlated pairs of lncRNA and sense transcript are in fact mutually exclusively expressed in different tissues or cell types in the perithecia, inhibiting each other.

Although genes found in sRNA-enriched loci are usually silenced by RNAi-dependent heterochromatin formation, some of the sRNA clusters involving lncRNAs, paradoxically, were transcriptionally active at the meiotic stage in *F. graminearum*. Since there are newly emerging tissues such as asci at the meiotic stage, it is conceivable that the silenced genes were derepressed by lncRNA expression in certain tissue types by yet-unknown mechanisms. A periodic gene activation and repression mechanism involving lncRNA and facultative heterochromatin formation has been documented in *N. crassa*. The expression of an lncRNA antisense to the circadian clock gene, *frequency*, promotes the sense transcription by generating a more transcriptionally permissive chromatin that has been silenced by sRNA-mediated heterochromatin formation (Li et al. 2015).

This study presents genome-wide characterization of lncRNAs during the fruiting body development. The transcriptome landscape including lncRNAs during the life cycle of *F. graminearum* provides fundamental genomic resources to the fungal community. The detailed molecular study of newly identified lncRNAs, with its established tools for rapid genetic analyses and ample genetic resources, will contribute to the understanding of how fungi utilize noncoding genomes for laying out their multicellular body plan.

## METHODS

### Data generation and processing

The *F. graminearum* genome assembly (Cuomo et al. 2007) and the Ensembl annotation version 32 (King et al. 2015) of a wild-type strain PH-1 (accessions: FGSC9075/NRRL31084) were used throughout this study. For total RNA extraction, synchronized fungal tissues were collected from carrot agar cultures at the previously defined developmental stages during perithecia formation (Trail and Common 2000; Hallen et al. 2007; Sikhakolli et al. 2012; Trail et al. 2017). For transcriptome data of spore germination stages, asexual spores (macroconidia) were spread on Bird agar medium (Metzenberg 2004) overlaid with a cellophane membrane, and sampled at the indicated spore germination stages (Supplemental Fig. S5). Strand-specific cDNA libraries were constructed from poly-A captured RNAs, using the KAPA Stranded RNA-Seq Library Preparation Kit (Kapa Biosystems, Wilmington, MA), and sequenced on the Illumina HiSeq 2500 platform (Illumina Inc., San Diego, CA) at the Michigan State University’s Research Technology Support Facility (https://rtsf.natsci.msu.edu/genomics). Following quality control for raw reads (Supplemental Methods), filtered reads were mapped to the repeat-masked genome using the HISAT2 program (v2.0.4; Kim et al. 2015a), and a genome-guided transcriptome assembly was performed using the StringTie program (v1.3.0) to generate *de novo* transcript annotations (Pertea et al. 2015).

### Differential expression and functional enrichment analyses

Read counts for gene loci were calculated using the *htseq-count* program (v0.6.1; Anders et al. 2015). On average, 87% of mapped reads were overlapped to exons in the *de novo* annotations. Gene expression levels in counts-per-million (CPM) value were computed and normalized by effective library size estimated by trimmed mean of M values, using the edgeR R package (v3.14.0; Robinson et al. 2010). Only genes with CPM greater than 1 in at least 3 samples were kept for further analyses (15,476 out of 20,459 gene loci). Then, differentially expressed (DE) genes showing greater than 4-fold difference at FDR 5% were identified between two successive developmental stages, using the limma R package (v3.28.21; Law et al. 2016). GO terms were assigned to the *de novo* annotations, using the Trinotate program (v3.0.1; Bryant et al. 2017). The list of GO terms was customized by adding several GO terms related to developmental processes to the GO Slim terms specific for fission yeast (Aslett and Wood 2006; Supplemental Table S1). Functional enrichment analyses for DE genes were performed using the GOseq R package (v1.24.0), including only those genes annotated by one or more GO terms (Young et al. 2010). To assess enrichment of GO terms, the Wallenius approximation (an extension of the hypergeometric distribution) and Benjamini–Hochberg method were used to calculate the FDR-corrected *P* value (Supplemental Table S1).

### Conserved lncRNA search

To search for conserved lncRNAs, the 547 lncRNAs were queried against the RNAcentral database version 5 (http://rnacentral.org), using the *nhmmer* program, which detects remote homologies, in the HMMER software (v3.1b2; Wheeler and Eddy 2013). Search hits with *E* < 10^−10^ were reported (Supplemental Table S3).

### Coexpression network analysis

The weighted gene correlation network analysis (WGCNA) R package (v1.51; Langfelder and Horvath 2008) was used to cluster lncRNAs by averaged RPKM values for developmental stages. The ‘*pickSoftThreshold*’ function was used to determine soft-thresholding power that measures the strength of correlation based on not just the direct correlation value of pairs of genes, but also the weighted correlations of all of their shared neighbors in the network space (Zhang and Horvath 2005). The soft-thresholding power 26 was selected, which is the lowest power for which the scale-free topology model fit index reaches 0.80. A range of treecut values were tested for cluster detection and the value was set to 0.18 (corresponding to correlation of 0.82). All other WGCNA parameters remained at their default settings.

### Identification of small RNA clusters

The sRNA-seq and degradome-seq data were obtained from NCBI GEO (GSE87835) and NCBI SRA (PRJNA348145), respectively (Son et al. 2017). In filamentous fungi, the size of a majority of sRNAs ranges from 17 to 27 nt with a strong 5′U preference (70–82%; Lee et al. 2010; Son et al. 2017). Thus, clusters of 17–27 nt-long 5′U-sRNA reads were detected across the genome, using the ShortStack program (v3.8.2; Shahid and Axtell 2014) with option arguments: ‘--pad 22’ and ‘--mincov 20’. Subsequently, the number of 5′U-sRNA reads aligned to different genomic features (*e.g.* coding regions) were counted for each sRNA cluster, using the *htseq-count* program (v0.8.0; Anders et al. 2015). The degradome-seq dataset was processed according to the previous study (Son et al. 2017).

### Expression correlation analysis

The expression value matrix for the 18 RNA-seq data (6 stages × 3 replicates) were rearranged by the BLIND program (Supplemental Fig. S2). Pearson’s correlation and the associated *P* value by Fisher’s exact test were calculated for the expression levels of the 300 sense mRNA-antisense lncRNA pairs, using an R script (Gaiti et al. 2015). For sense mRNAs only, that showed positive correlation with antisense lncRNAs, we performed functional enrichment analyses, using the GOseq R package (v1.24.0; Young et al. 2010), as did for DE genes between two successive developmental stages (see above).

### XUT identification

Generation and confirmation of gene-deletion mutants and RNA-seq methodology for Δ*xrn1* were described in Supplemental methods. For Δ*xrn1* RNA-seq data, transcripts were independently assembled and merged to the *de novo* annotations, using the ‘*merge*’ function in the StringTie program (v1.3.0; Pertea et al. 2015), to incorporate novel transcripts that were only expressed in Δ*xrn1* due to loss of NMD activity. Transcript abundance is globally affected by *XRN1* deletion in budding yeast, however the expression of ribosomal protein genes shows only slight increases in Δ*xrn1* and remains constant after lithium treatment that inhibits 5′→3’ exonuclease activities (Dichtl et al. 1997; van Dijk et al. 2011). Thus, before XUT identification, expression values were normalized in such a way that ribosomal protein genes are expressed at the same levels in WT and Δ*xrn1* by multiplying 0.66 and 1.96 to RPKM values of WT at S0 and S4, respectively (Supplemental Fig. S11). Differential expression analyses were performed, using the ‘*Stattest*’ function with an option argument: ‘libadjust=FALSE’, in the Ballgown R package (v2.8.4; Frazee et al. 2015). The DE transcripts showing 3-fold increase in Δ*xrn1* were identified as XUTs (5% FDR). We excluded transcripts with relatively low expression levels in Δ*xrn1* (RPKM < 3.0) from the putative XUTs to reduce possible false positives.

### DATA ACCESS

The RNA-seq data generated in the present work have been deposited in NCBI’s Gene Expression Omnibus (https://www.ncbi.nlm.nih.gov/geo), and are accessible through GEO series accession number GSE109095 that is composed of the two datasets, the sexual stage and Δ*xrn1* dataset (GSE109094) and the spore germination stage dataset (GSE109088).

## ACKNOWLEDGMENTS

We thank Brad Cavinder, Kevin Childs, and Nicholas Panchy (Michigan State Univ., MI, USA) for helpful discussion on analyzing data and for providing Perl and Python scripts. This work was supported by the National Science Foundation (Grant IOS-1456482 to F.T.) and Michigan AgBioResearch (F.T.). The funders had no role in study design, data collection and analysis, decision to publish, or preparation of the manuscript.

This work was supported by the Program Area: Plant Health and Production and Plant Products [Grant no. 2015-67013-22932 to F.T.] from the USDA National Institute of Food and Agriculture. Any opinions, findings, conclusions, or recommendations expressed in this publication are those of the authors and do not necessarily reflect the view of the U.S. Department of Agriculture.

## Author contributions

W.K. and F.T. conceived and designed the experiments; W.K. and C.M-R. performed the experiments; W.K., J.W., J.P.T. and F.T. analyzed the data; W.K. wrote the paper; and J.P.T. and F.T. revised the paper.

